# LSD1/KDM1A Maintains Genome-wide Homeostasis of Transcriptional Enhancers

**DOI:** 10.1101/146357

**Authors:** Saurabh Agarwal, Patricia Marie Garay, Robert Scott Porter, Emily Brookes, Yumie Murata-Nakamura, Todd S. Macfarlan, Bing Ren, Shigeki Iwase

**Author notes:** Correspondence (S.I.), (B.R.).

## Abstract

Transcriptional enhancers enable exquisite spatiotemporal control of gene expression in metazoans. Enrichment of mono-methylation of histone H3 lysine 4 (H3K4me1) is a major chromatin signature that distinguishes enhancers from gene promoters. Lysine Specific Demethylase 1 (LSD1, aka KDM1A), an enzyme specific for demethylating H3K4me2/me1, has been shown to “decommission” stem cell enhancers during the differentiation of mouse embryonic stem cells (mESC). However, the roles of LSD1 in undifferentiated mESC remain obscure. Here, we show that LSD1 occupies a large fraction of enhancers (63%) that are primed with binding of transcription factors (TFs) and H3K4me1 in mESC. In contrast, LSD1 is largely absent at latent enhancers, which are not yet primed by TF binding. Unexpectedly, LSD1 levels at enhancers exhibited a clear positive correlation with its substrate, H3K4me2 and enhancer activity. These enhancers gain additional H3K4 methylation upon the loss of LSD1 in mESC. The aberrant increase in H3K4me at enhancers was accompanied with increases in enhancer H3K27 acetylation and expression of enhancer RNAs (eRNAs) and their target genes. In post-mitotic neurons, loss of LSD1 resulted in premature activation of enhancers and genes that are normally induced after neuronal activation. These results demonstrate that LSD1 is a versatile suppressor of primed enhancers, and is involved in homeostasis of enhancer activity.

## Introduction

Transcriptional enhancers were discovered as potent gene regulatory elements that act independently of the distance and orientation to the target promoters ^1, 2^. A broad range of physiological and developmental processes rely on coordinated actions of transcriptional enhancers to achieve cell type-specific and temporally-controlled gene expression ^3, 4^. Numerous non-coding variants associated with a variety of human traits have been observed at enhancers, implicating their importance in normal physiology and disease pathogenesis ^5-7^.

Recent studies have begun to reveal how the life cycle of enhancers progresses to induce gene expression changes during development ^8^. Genome-wide discovery of thousands of potential enhancer elements has been facilitated by profiling 1) binding of pioneer transcription factors (TFs), 2) chromatin accessibility as measured by hypersensitivity to DNase I, and 3) patterns of histone modifications (reviewed in ^9^). Monomethylation of histone H3 lysine 4 (H3K4me1) can distinguish enhancers from promoters, which in contrast are modified with H3K4me3 ^10, 11^. In response to various environmental and developmental cues, TFs bind to specific DNA elements ^12, 13^, and subsequent recruitment of methyltransferases KMT2C and KMT2D (aka MLL3 and MLL4, respectively) leads to H3K4me1 at enhancers ^14-16^.

Once installation of H3K4me1 “primes” enhancers, they can become either “active” or “poised”, depending on the acetylation or tri-methylation of H3 lysine 27 (H3K27ac or H3K27me3) ^17, 18^, respectively. A recent report showed that active enhancers can be negatively regulated by RACK7-mediated recruitment of KDM5C, an H3K4me3/2 demethylase ^19^. When genes need to be turned off, e.g. pluripotency genes during differentiation of stem cell, their enhancers undergo “decommissioning” by LSD1-mediated removal of the priming mark, H3K4me1 ^20^. Notably, “latent” enhancers, i.e. DNA elements that lack TF binding or H3K4me1, can gain enhancer-like marks in response to extra-cellular stimuli and promote gene expression in fully-differentiated macrophages ^21^. These findings highlight the dynamic H3K4 methylation of an enhancer during its life cycle.

Two H3K4 demethylases, LSD1 and KDM5C, have been shown to play important roles in regulation of enhancers ^19, 20, 22^. While KDM5C reverses H3K4me3/2 leaving H3K4me1 intact ^23, 24^, LSD1 can demethylate only H3K4me2/1 ^25^. The distinct substrate specificities raise a possibility that these two H3K4 demethylases may cooperate to generate and/or maintain the balance of H3K4me landscape at different classes of enhancers. For example, absence of H3K4 methylation at latent enhancers could potentially be attributed to LSD1-mediated demethylation of H3K4me1. Besides the decommissioning of stem cell genes and enhancers during differentiation, LSD1 has also been shown to repress developmental genes ^26, 27^ and retrotransposons ^28^ in ES cells. However, it remains unclear whether LSD1 and KDM5C play any role at other classes of enhancers.

In the present study, we demonstrate that in addition to active enhancers, LSD1 also occupies poised enhancers, some of which are quickly activated (inducible enhancers) upon differentiation of mESC or depolarization of post-mitotic neurons. Interestingly, LSD1 does not bind to latent enhancers, e.g. neuron-specific enhancers that are unprimed in mESC, and LSD1 occupancy shows a clear positive correlation with its substrate, H3K4me2. Loss of LSD1, but not KDM5C, leads to a global upregulation of enhancer RNAs accompanied with increased H3K4 methylation and H3K27ac at active, poised, and inducible enhancers, and their target genes. These results indicate that LSD1 is a pervasive suppressor of primed enhancers, involved in negative-feedback mechanisms to maintain homeostasis of histone-modification landscapes at enhancers.

## Results

### LSD1 occupies a large fraction of primed enhancers

To study the role of LSD1 in regulation of enhancers and gene expression, we first examined the genome-wide distribution of LSD1 at various regulatory elements. We analyzed the previously published ChIP-Seq datasets of LSD1 ^20^, p300 ^29, 30^, CTCF ^29, 31^, DNase-Hypersensitivity (DHS) ^32^ and other histone modifications (see Supplementary Table 1). p300, a histone acetyltransferase and a transcriptional coactivator, has been shown to occupy both promoters and enhancers ^10, 33^, whereas CTCF binding sites anchor chromatin loops ^34^ and insulating domains ^35-37^. By examining the overlaps of binding sites of LSD1 (109,541, *q* < 0.05), p300 (86,426, *q* < 0.01), CTCF (58,899, *p* < 10^−12^) and DHS sites (299,799, *q* < 0.05), we found that 1) a majority of p300 binding sites (70.5%, Figure 1a) were co-occupied by LSD1, 2) in contrast, only 14.7% of non-p300 CTCF-binding sites were occupied by LSD1, and 3) most of the LSD1 binding sites (86%) showed an overlap (± 250 bases) with DHS sites. The higher degree of overlap of LSD1 with p300 compared to CTCF-only sites was observed at promoter, genic, and intergenic regions (Supplementary Figure 1). These observations indicate that LSD1 occupies a large fraction of primed enhancers.

**Figure 1.**
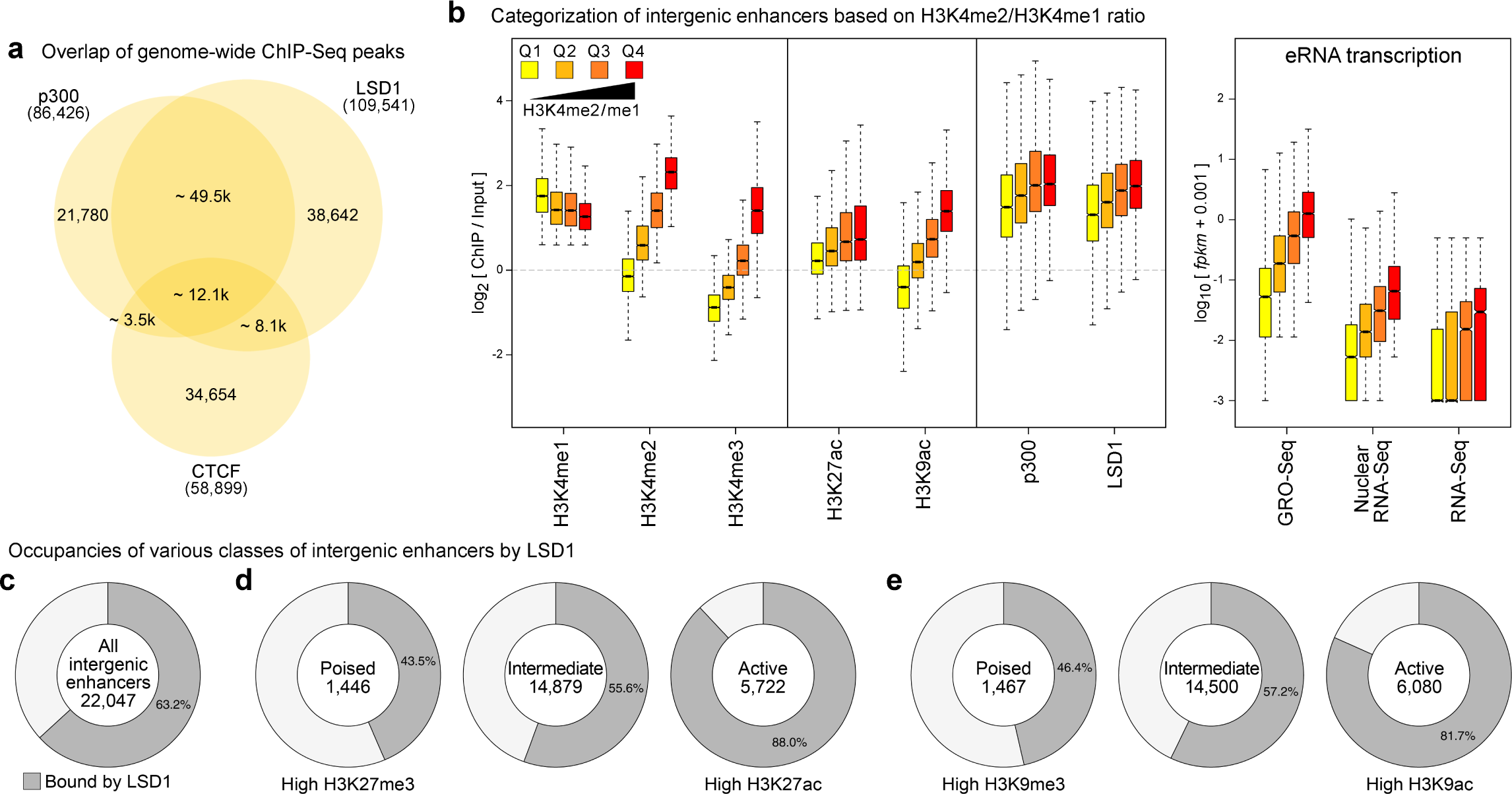
LSD1 occupies a large fraction of primed enhancers in mES cells. (**a**) Overlap of binding sites of p300, CTCF, and LSD1 in mESC. (**b**) Intergenic enhancers were divided into quartiles (Q1-Q4) based on the enrichment of H3K4me2 relative to H3K4me1 (left panel). Boxplots of enrichment of indicated histone modifications, LSD1, and p300, as measured by ChIP-Seq, and eRNA levels (GRO-Seq, Nuclear RNA-Seq, and RNA-Seq) at each quartile of intergenic enhancers. Levels of LSD1 show positive correlations with increases in H3K4me2 and eRNA expression from Q1 to Q4. (**c**) The percentage of intergenic enhancers with LSD1 peaks. (**d, e**) LSD1 occupancy at active, poised, and intermediate enhancers classified based on enrichment of either trimethylation or acetylation of H3K27 (d) or H3K9 (e). LSD1 occupancy at enhancers increases with higher activity. In all figures, the bottom and top boxes signify the second and third quartiles, respectively, and the middle band represents the median of the population. Whiskers represent 1.5 times the inter-quartile range (IQR) and the notch represents the 95% confidence interval of the median.

Next, we sought to identify regulatory elements that could potentially act as enhancers. Previous studies have utilized a high H3K4me1:me3 ratio, either alone ^10, 17^ or in conjunction with TF binding ^38, 39^, DHS or binding by CBP/p300 ^40, 41^, to distinguish enhancers from promoters in a given cell-type. H3K4me2 is observed at both promoters and enhancers and has been shown to be a signature to predict enhancers ^38^. We therefore included H3K4me2 data to increase the sensitivity and precision of enhancer mapping. Enhancers also differ from promoters in that promoters are associated with stable transcripts ^42^ while active enhancers are associated with expression of enhancer RNA transcripts (eRNAs) ^42-44^, which are short-lived due to exosome-mediated degradation ^44-46^. This degradation of nascent transcripts from enhancers results in a very low, albeit, detectable levels of eRNAs in RNA-Seq (Figure 1b, right panel). Global Run-On, an *in vitro* assay, followed by high-throughput sequencing (GRO-Seq) ^47^ enables a sensitive and quantitative evaluation of transcriptionally-engaged RNA polymerase molecules. GRO-Seq, thus, serves as an indirect measure of nascent transcription at promoters and enhancers, irrespective of the subsequent stability of the transcripts ^42^. Therefore, we employed a high ratio of GRO-Seq:RNA-Seq signals to further refine the prediction of enhancers in mESC. We focused on only intergenic enhancers as we found it difficult to differentiate the eRNAs from gene-coding and promoter-upstream ^46^ transcripts. In summary, intergenic enhancers were defined as ± 500 base regions around p300/DHS summits with i) H3K4me1 enrichment (*rpkm* ≥ 1 and ChIP:Input > 1.5), ii) H3K4me3 lower than either H3K4me1 or H3K4me2, iii) a low rate of transcription (RNA-Seq *fpkm* < 0.5), iv) a GRO-Seq:RNA-Seq ratio > 5, and v) a high average mappability to exclude repetitive regions. This pipeline predicted a total of 22,047 intergenic enhancers in mESC (Supplementary Table 2).

LSD1 has been shown to occupy enhancers in various cell types ^20, 48, 49^. However, the genome-wide relationship between LSD1 binding and chromatin states at enhancers, such as histone-modification landscapes and eRNA transcription, remains unclear. To address this issue, we first subdivided the 22,047 predicted intergenic enhancers into quartiles (Q1-Q4, Figure 1b) based on the enrichment of H3K4me2 relative to H3K4me1. Similar to a previous observation in K562 cells ^42^, we noted a positive correlation between eRNA levels, measured by GRO-Seq, Nuclear RNA-Seq or RNA-Seq, and H3K4me2 levels (Figure 1b). Acetylations of H3K9 and H3K27 and eRNA expression have been established as signatures of active enhancers ^9, 17, 18^. Consistently, we also observed that enhancers with higher transcription levels displayed higher acetylation levels of H3K9 and H3K27 relative to trimethylation (Supplementary Figure 2).

We then determined the extent of LSD1 binding across various enhancer classes and found that a large fraction (63.2%) of the predicted 22,047 enhancers is bound by LSD1 (Figure 1c). Surprisingly, we found that LSD1 occupancy at enhancers increased with increasing levels of H3K4me2 or increasing levels of GRO-Seq signals (Supplementary Figure. 3 and 4a). We then calculated the correlation coefficients between LSD1 levels and levels of various histone modifications at promoter distal regions, i.e. excluding TSS ± 1500 bases. Compared to H3K4me1 (*r* = 0.5745) and H3K4me3 (*r* = 0.486), we found that LSD1 levels showed the highest correlations with its primary substrate, H3K4me2 (*r* = 0.627, Supplementary Figure 4b), and H3K27ac (*r* = 0.604), a marker for enhancer activity. In contrast, H3K27me3 and H3K9me3 were inversely correlated with LSD1 levels (Supplementary Figure 4b). We also classified 22,047 intergenic enhancers into poised, active, and intermediate enhancers based on their H3K27me3 and H3K27ac levels, according to the previous report ^18^. We found that LSD1 occupies substantial fractions of each of the three enhancer classes with increased occupancy of active enhancers compared to other classes (Figure 1d). Similar patterns were observed when we used the levels of H3K9me3 and H3K9ac to classify enhancers as described previously ^41^(Figure 1e). These results indicate that LSD1 binds to multiple enhancer classes with positive correlations to H3K4me2, H3K27/K9 acetylation and eRNA levels.

### LSD1 rarely binds to cell-type specific “latent” enhancers

Higher LSD1 occupancy at more active enhancers may contradict with the LSD1’s classical role as a transcriptional repressor of neuron-specific genes in non-neuronal tissues ^25, 50^. RE1-Silencing Transcription factor (REST) is known to be expressed in non-neuronal cells with the role of repression of neuronal genes in these cell-types through the Corepressor of REST(CoREST) complex ^50^. We analyzed the previously published REST ChIP-Seq dataset ^20^ and found that only 1.31% of predicted enhancers (288 out of 22,047) were bound by REST in mESC. However, a majority (91%) of these REST-positive enhancers were bound by LSD1. These data suggest that REST/LSD1-mediated suppression is not the primary mechanism of enhancer regulation in mESC. Enhancers that are inert, free of TFs, and thus, insensitive to DNase I, are referred to as “latent” enhancers in a given cell type ^21^. The low occupancy of primed enhancers by REST/LSD1 prompted us to test if LSD1 contributes to regulation of latent enhancers. We first looked at the *mir290* cluster, which is specifically expressed in mESC and early developmental stages ^51^. In mESC, several enhancers upstream of its promoter, show high DHS, p300 binding and high GRO-Seq signals accompanied with strong LSD1 binding events (Figure 2a). These *mir290* enhancers lack brain-derived DHS, GRO-Seq signal and H3K4 methylation in cortical neurons (CN), demonstrating that these enhancers are primed in mESC but are latent in CN. LSD1 ChIP-Seq data from neural stem cells (NSC) ^52^, however, showed a lack of LSD1 binding at these latent enhancers. Conversely, two enhancers upstream of *Npas4* (marked with asterisks, Figure 2b), a gene predominantly expressed in the brain, showed brain-specific DHS and LSD1 occupancy, GRO-Seq signals and high H3K4me1, specifically in the neuronal cell types (NSC or CN). In mESC, LSD1 is absent at these brain-specific DHS sites upstream of the *Npas4* promoter (Figure 2b). These two examples suggest that LSD1 could primarily be recruited to primed enhancers in a given tissue in a TF-binding dependent manner.

**Figure 2.**
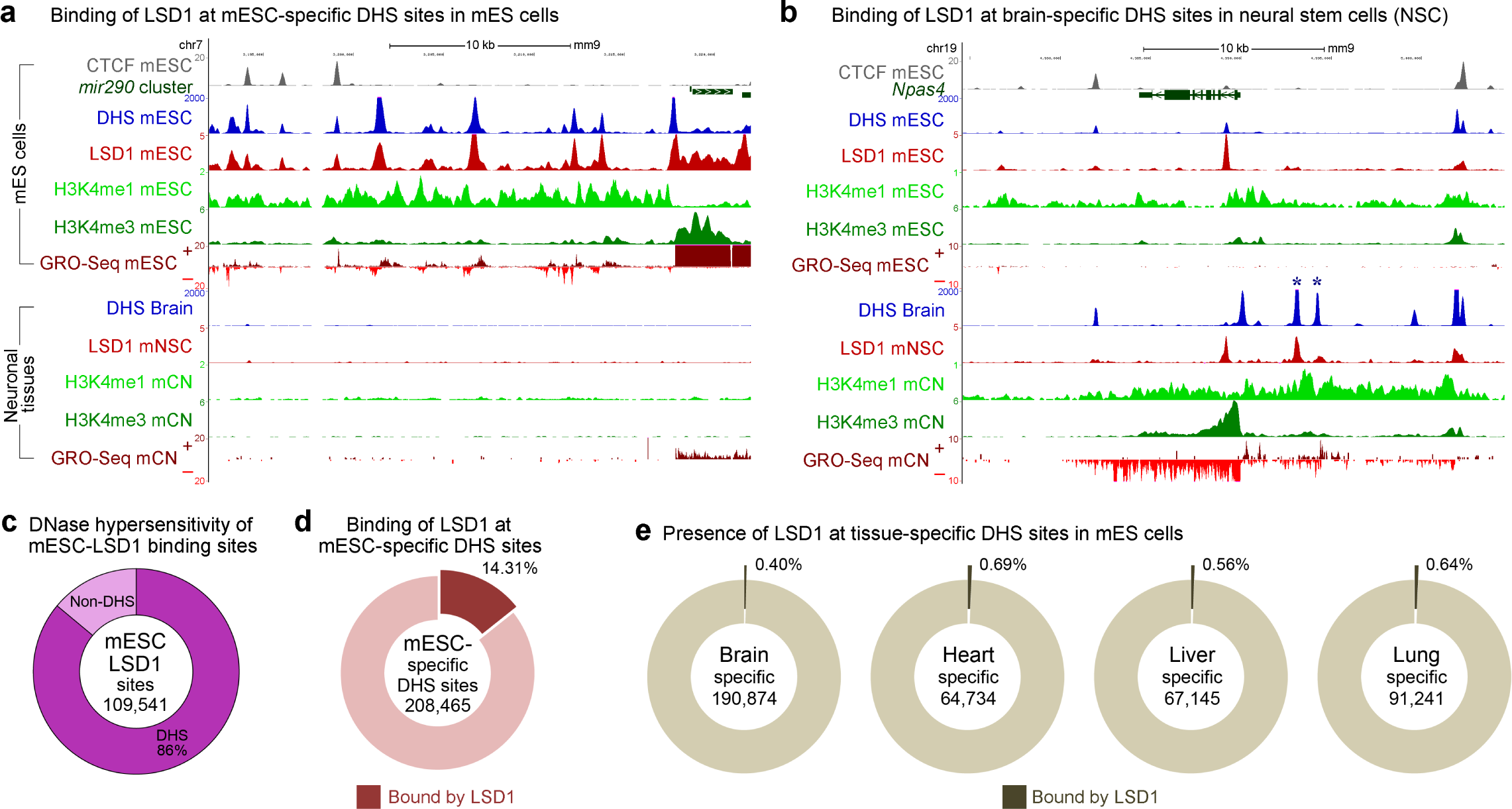
LSD1 rarely binds to cell-type specific “latent” enhancers. (**a**) UCSC genome browser snapshot of the *mir290* locus. LSD1 ChIP-Seq peaks in mESC coincide with enhancer signatures, including DHS, H3K4me1, and divergent GRO-Seq signals, upstream of the *mir290* promoter in mESC but not in neuronal cells. (**b**) LSD1 binding at the *Npas4* locus in mESC and NSC. LSD1 is present at the two brain-specific DHS sites (marked by asterisks) in neuronal cells but not in mESC. Note that the DHS sites common in adult brain and mESC are occupied by LSD1 in both mESC and NSC. NSC: Neural stem cells, CN: Cortical neurons. CTCF, DHS and LSD1 tracks were generated from previously published datasets. (**c**) Fraction of LSD1 peaks overlapping with mESC hotspots. (**d**) Fraction of mESC-specific hotspots overlapping with mESC-specific LSD1 peaks. (**e**) Fractions of tissue-specific hotspots overlapping with LSD1 peaks with no mESC-derived DHS.

To ascertain this specificity of LSD1 recruitment to primed enhancers on a genome-wide scale, we sought to identify genomic elements that are latent in mESC but are primed in other cell types. We identified DHS sites from mESC (398,675, *q* < 0.01) and four additional mouse tissues, including adult brain (415,400), heart (320,416), liver (207,046), and lung (358,575), using Hotspot (v4.1) ^53^. Similar to our earlier observation (Supplementary Figure 1), we found that most of mESC LSD1-binding sites (86%) overlapped with mESC hotspots (Figure 2c). Next, we performed an intersection of hotspots from the five tissues. This resulted in tens of thousands of hotspots, which could potentially act as tissue-specific enhancers in a given tissue and latent in others (Figure 2d, e). Motif analysis on promoter-distal hotspots revealed that these tissue-specific hotspots are indeed enriched with binding sites for lineage-specific TFs (Supplementary Figure 5). In agreement with the *mir290* and *Npas4* loci, mESC LSD1 binding sites showed negligible overlaps with tissue-specific hotspots (0.40-0.69%), whereas 14.31% of mESC-specific hotspots were bound by LSD1 (Figure 2d, e). Based on these data, we concluded that LSD1 is predominantly recruited to primed enhancers and is not actively involved in maintaining inactivity of latent enhancers in mESC.

### Loss of LSD1 results in a genome-wide increase in enhancer H3K4 methylation and H3K27 acetylation

The unexpected positive correlation between LSD1 binding and enhancer H3K4me2 levels raises a possibility that LSD1 may not be demethylating H3K4me2 at enhancers. LSD1 has been implicated in demethylation of H3K9 instead of H3K4 when it binds androgen receptors ^54, 55^, though a recent study reported otherwise ^56^. Phosphorylation of H3T6 appears to interfere with LSD1-mediated H3K4 demethylation ^57^. Alternatively, the positive correlation may reflect a negative feedback mechanism, in which LSD1 searches for and binds to genomic regions with high H3K4me2 levels and reverses this modification to regulate optimal enhancer activity.

To test whether LSD1 is involved in maintaining precise levels of H3K4 methylation at enhancers, we investigated the previously generated mESC line that lacks LSD1 due to the insertion of a gene-trap cassette (*Lsd1*-GT) ^28^. Western blot analysis of mESC carrying either wild-type (WT) *Lsd1* or *Lsd1*-GT did not show any detectable differences in total H3K4me1, H3K4me2, H3K4me3 or H3K27ac levels (Supplementary Figure 6). We then performed ChIP-Seq to measure H3K4me levels across the genomes of these two mESC lines. Since LSD1 is known to associate with multiple HDAC-containing co-repressor complexes, including the CoREST ^50, 58^ and NuRD ^59^ complexes, we also included H3K27ac and HDAC1 in our ChIP-Seq analysis.

Genome-wide localization analysis (ChIP-Seq) profiles, which reflect the spatial distribution of these marks, looked highly similar between the two genotypes at most of the loci. Upon the loss of LSD1, however, H3K4 methylations displayed statistically-significant increases at active, poised, and intermediate LSD1-target enhancers, which were accompanied with conspicuous increases in H3K27ac (Figure 3a, Supplementary Figure 7a). Similar changes were also observed at enhancers that showed a significant increase in eRNA expression (see next section) in the *Lsd1-*GT mESC (Supplementary Figure 7). Interestingly, HDAC1 levels did not change significantly at poised enhancers, while active or poised enhancers showed a small but significant increase in HDAC1 binding (Figure 3a), which could be attributed to either experimental variations or unknown mechanisms to compensate for the loss of LSD1. The inability of HDAC1 to remove H3K27ac in *Lsd1*-GT cells is consistent with the previous observations that HDAC activity is negatively influenced by the presence of H3K4me ^27, 60^. Representative genes *Pou5f1* (Figure 3b) and *Cbln4* (Figure 3c), which are normally active or poised in undifferentiated mESC, respectively, showed relatively higher H3K4me and H3K27ac at both promoters and enhancers in *Lsd1-*GT mESC. We also observed a concomitant increase in both promoter- and enhancer-associated GRO-Seq signals at these loci in *Lsd1-*GT mESC (Figure 3b, c). The increases in H3K27ac and nascent transcription at these poised and intermediate enhancers upon the loss of LSD1 suggest a shift in their identity towards active enhancers. These results indicate that LSD1 functions as an H3K4 demethylase at enhancers, and is required for maintenance of optimal H3K4me and H3K27ac levels in mES cells.

**Figure 3.**
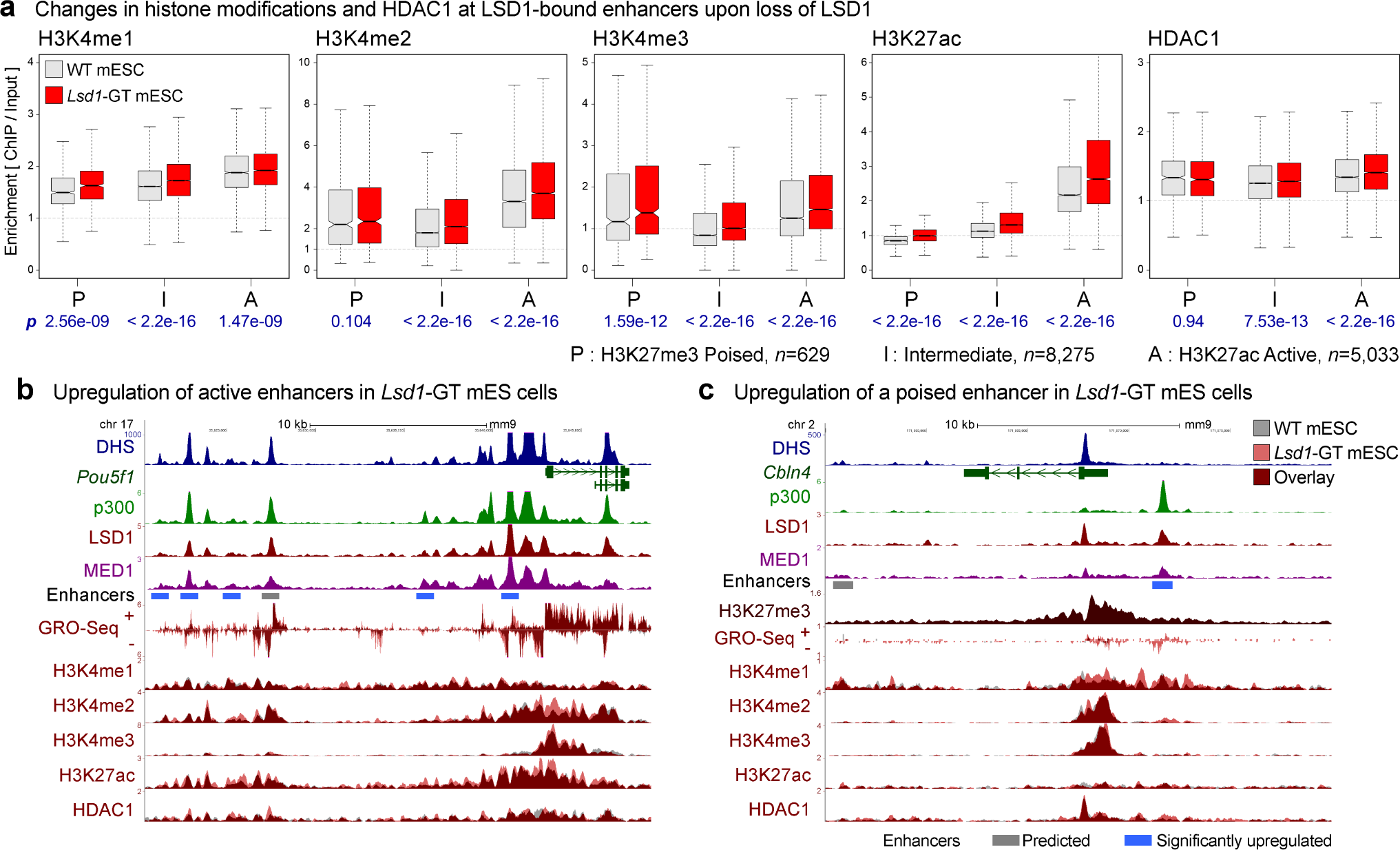
Loss of LSD1 results in increases in H3K4 methylation and H3K27 acetylation at enhancers. (**a**) H3K4me1, H3K4me2, H3K4me3, H3K27ac and HDAC1 levels on LSD1-bound enhancers in WT (gray boxes) and *Lsd1*-GT mESC (red boxes). Enhancers were classified into poised (P), intermediate (I), and active (A) enhancers based on the enrichment of either H3K27ac or H3K27me3. Geometric mean of ChIP:Input ratios from the two independent ChIP-Seq replicates are shown. *P*-values (*p*) from Wilcoxon signed-rank tests on differences, log2(*Lsd1*-GT/WT), are denoted in blue beneath each panel. *n* indicates the number of enhancers in each category. (**b**) Dysregulation of active enhancers at the *Pou5f1* (aka *Oct4*) locus. A cluster of enhancers is co-occupied by p300 and LSD1. Some of the individual enhancers show an increase in H3K4me2, H3K27ac, and GRO-Seq signals in *Lsd1*-GT mESC (red) compared to WT mESC (gray). (**c**) Misregulation of a poised enhancer (red bar) at the *Cbln4* locus. This locus is decorated with a broad H3K27me3 domain, and shows elevation in H3K4me1, H3K4me2, and GRO-Seq signals upon the loss of LSD1. Gray bar: Predicted enhancer. Blue bar: significantly-upregulated enhancer in *Lsd1*-GT mESC compared to WT mESC based on changes in GRO-Seq signal (See Figure 4).

### Loss of LSD1 but not KDM5C results in aberrant activation of transcriptional enhancers

In the above ChIP-Seq study, we found a genome-wide elevation of all three H3K4me statuses, including H3K4me3, at LSD1-target enhancers in *Lsd1*-GT mESC (Figure 3a, Supplementary Figure 7a). This increase of H3K4me3 at enhancers cannot be explained directly by the loss of LSD1, as LSD1 is incapable of demethylating H3K4me3 ^25^; therefore, one or more H3K4me3 demethylases might be involved in maintaining low levels of H3K4me3 at enhancers. LSD1 and KDM5C, an H3K4me3/me2 demethylase ^23^, have been previously shown to be in the same complex ^19^ and that KDM5C suppresses over-activation of active enhancers in breast cancer cells ^19^. To elucidate the interrelationship between LSD1 and KDM5C in suppression of enhancer activity, we first performed KDM5C ChIP-Seq in mESC and identified 113,166 KDM5C binding sites (MACS2, *q* < 0.05). Most of the 22,047 predicted intergenic enhancers (78.3%) were bound by either LSD1 or KDM5C and 52.1% of total were bound by both (Figure 4a).

**Figure 4.**
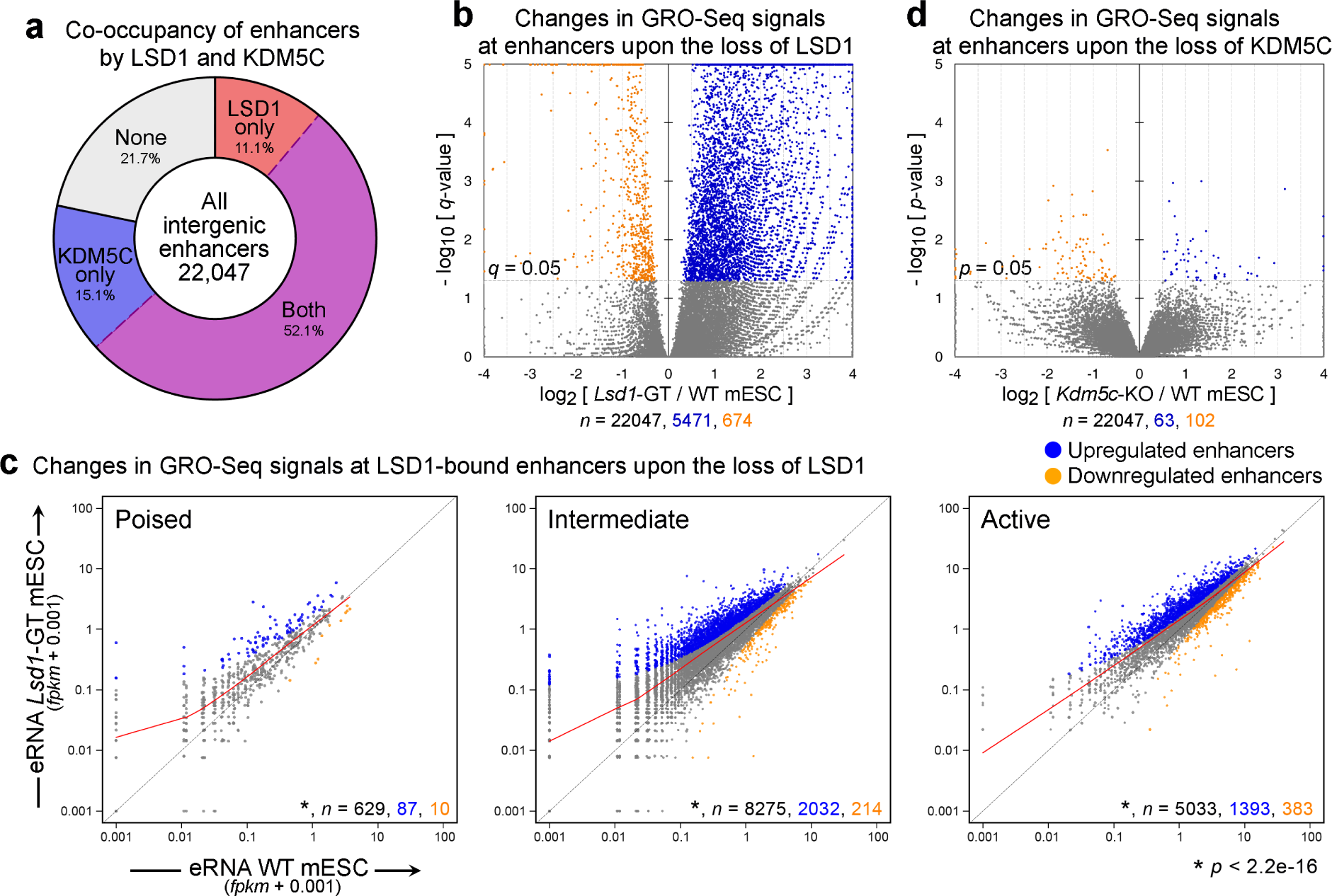
Loss of LSD1 but not KDM5C results in aberrant activation of enhancers. (**a**) Fractions of intergenic enhancers bound by LSD1 and/or KDM5C in mESC. (**b, d**) Volcano plots of GRO-Seq signals at all intergenic enhancers from DESeq analysis. While the loss of LSD1 resulted in a large-scale increase in GRO-Seq signals at enhancers, deletion of KDM5C had a minimal impact. X-axis and Y-axis indicate the log2 fold-change and significance, respectively of differential expression in WT and mutant mES cell lines. (**c**) Scatter plots of GRO-Seq levels at poised, intermediate, and active enhancers classified of the basis of enrichment of either H3K27me3 or H3K27ac. Significantly-upregulated and -downregulated enhancers (*q* < 0.05, DESeq) are shown in blue and orange, respectively. Red curve indicates the LOWESS curve for each class of enhancers. Total number (*n*) of all, significantly-upregulated, and -downregulated enhancers in each group are indicated in black, blue, and orange, respectively. Each class of enhancers shows a significant upregulation (*p* < 2.2e-16, Wilcoxon signed-rank test) in *Lsd1*-GT mESC compared to WT mESC.

We generated *Kdm5c*-knockout (KO) mESC by transfecting a *Cre*-expression plasmid into the mESC harboring the floxed exons 11 and 12, which encode the catalytic JmjC domain ^61, 62^, and confirmed the loss of KDM5C (Supplementary Figure 8). We then asked if the loss of either LSD1 or KDM5C leads to aberrant enhancer activity by quantifying changes in GRO-Seq signals at enhancers. To identify misregulated enhancers, we calculated the number of GRO-Seq reads mapping within ± 500 bases of the center of the predicted enhancers and normalized them against 199,209 p300/DHS sites across the whole genome using DESeq ^63^. Upon the loss of LSD1, a large fraction (24.8%, 5,471) of total intergenic enhancers showed a significant elevation in associated GRO-Seq transcripts, while a small number 674 (3.06%) displayed a reduced activity with a stringent cutoff of *q* < 0.05 (Figure 4b). Next, we tested if this elevation of GRO-Seq signals is specific to poised, intermediate or active enhancers. We found that all three enhancer classes showed a significant increase in associated nascent transcripts (*p* < 2.2e-16, Wilcoxon signed-rank test, Figure 4c), indicating that LSD1 is required for genome-wide suppression of aberrant enhancer activities.

Using the same DESeq cutoff, however, we were not able to identify any misregulated enhancers in *Kdm5c*-KO mESC. After relaxing the cutoff to *p* < 0.05, we could identify only 63 upregulated and 102 downregulated enhancers upon the loss of KDM5C (Figure 4d). To confirm that these observations were not dependent on differences in sequencing depths or inter-replicate variability, equal number of reads were randomly selected from each GRO-Seq sample and pairwise DESeq comparisons between individual replicates of either genotypes were repeated. Thus, in contrast to the crucial role of LSD1, KDM5C is largely dispensable for enhancer suppression in mESC.

Since GRO-Seq is an *in vitro* transcription assay, we sought to validate this global upregulation of enhancers upon LSD1 depletion in mESC under physiological conditions by sequencing total cellular RNAs (RNA-Seq) and nuclear RNAs (Nuclear RNA-Seq). Either RNA-Seq or Nuclear RNA-Seq could not provide sufficiently high eRNA signals to call differentially-expressed enhancers likely due to the aforementioned exosome-mediated degradation of eRNAs. However, when we evaluated eRNA levels at all the intergenic enhancers as a group, RNA-Seq and Nuclear RNA-Seq corroborated our GRO-Seq results (Supplementary Figure 9). These data demonstrate that LSD1, but not KDM5C, is required for suppression of aberrant enhancer activities in mESC.

### Aberrant changes in enhancer activity are associated with misregulation of physically-interacting genes

The standard approach to gauge the influence of enhancer misregulation on gene expression has been to quantify changes in expression of genes that are located in proximity to the enhancers of interest. However, recent advances in genome-wide profiling of chromatin interactions ^64-66^ have paved the way for a more precise determination of enhancer-promoter interactions. To identify genes that physically interact with our set of predicted enhancers, we utilized the recently-published “HiCap” data set, which is a high-resolution map of promoter-anchored chromatin interactions in mESC ^67^. For instance, our enhancer prediction identified a ~ 6 kb-wide enhancer cluster downstream of *Dusp5* and the analysis of HiCap data revealed that one of the three individual enhancers within the cluster appears to interact with the *Dusp5* promoter (Figure 5a). This enhancer cluster was significantly upregulated in *Lsd1*-GT mESC and showed a pronounced increase in H3K4me2 and H3K27ac levels, and *Dusp5* transcription (Figure 5a). A concomitant misregulation of enhancers and the interacting gene was also observed for the aforementioned *mir290* cluster (Supplementary Figure 10a).

**Figure 5.**
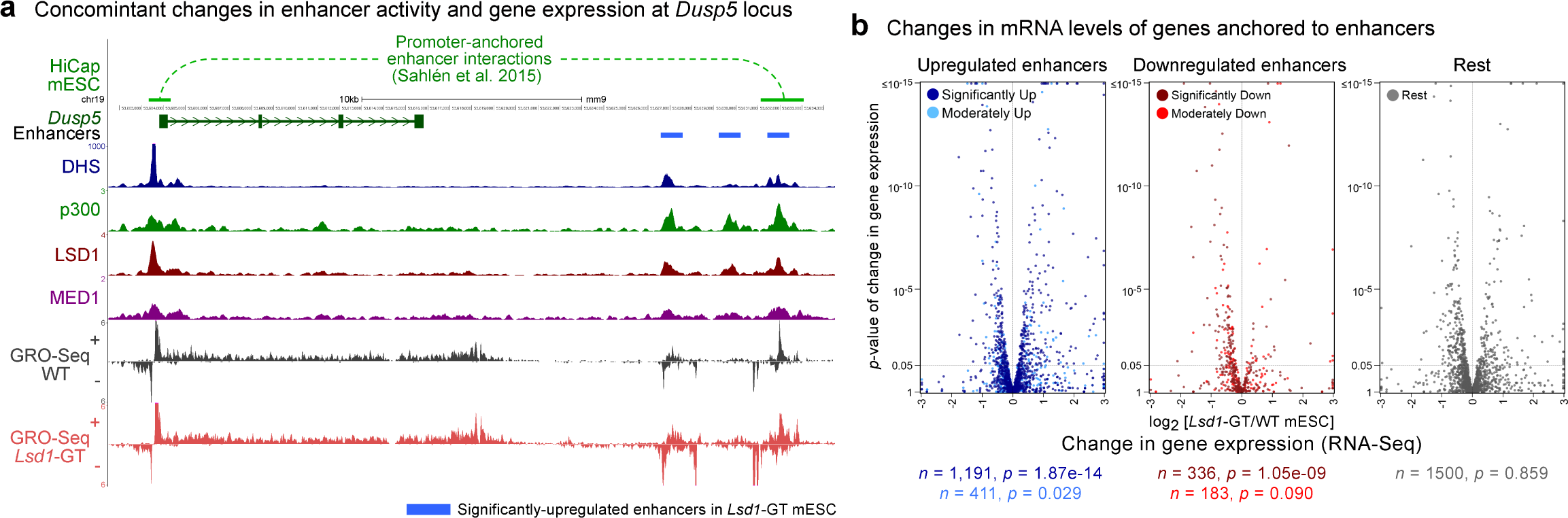
Aberrant changes in enhancer activity are associated with misregulation of physically-interacting genes. (**a**) An example of long-range promoter-enhancer interactions (top track) obtained from the mESC HiCap data set ^67^ at the *Dusp5* locus. One of the three significantly-upregulated enhancers (blue bars) interacts with the *Dusp5* promoter. Upon the loss of LSD1, the gene and the enhancers show upregulation of H3K4me2, H3K27ac and GRO-Seq signals in *Lsd1*-GT mESC (red) compared to WT mESC (gray). (**b**) Volcano plots of changes in mRNA levels (RNA-Seq) of genes that physically interact with misregulated enhancers. Based on changes in enhancer-associated GRO-Seq signals upon the loss of LSD1, enhancers were subdivided as significantly up (*q* < 0.05, DESeq), significantly down, moderately up (0.05 ≤ *q* < 0.25), moderately down, and the rest. When multiple enhancers showed interactions with a single promoter, assignment of the genes to each enhancer subgroup was prioritized in the aforementioned order. Total number of associated genes (*n*) and *p*-values (*p*) from Wilcoxon signed-rank tests on differences between mRNA levels in *Lsd1*-GT and WT mESC are shown beneath each panel. Note that more genes anchored to upregulated enhancers are upregulated compared to genes that interact with downregulated enhancers.

To evaluate the genome-wide impact of enhancer misregulation on gene expression, we first categorized enhancers based on the statistical significance of their differential expression in *Lsd1*-GT mESC: significantly misregulated (*q* < 0.05) and moderately misregulated (0.05 ≤ *q* < 0.25) enhancers from our GRO-Seq analysis. We then retrieved the physically-interacting promoters from the HiCap data, and plotted the changes in mRNA levels (RNA-Seq, Figure 5b) or rates of nascent transcription (GRO-Seq, Supplementary Figure 10b). We observed a general trend that genes associated with upregulated enhancers showed an increased expression and *vice versa*, and the genes that are anchored to unaffected enhancers did not exhibit any significant changes in expression in *Lsd1*-GT mESC (Figure 5b, Supplementary Figure 10b, c). Importantly, for each category, the magnitude and the statistical significance of median change in gene expression correlated positively with those of changes in enhancer activity (Supplementary Figure 10c). Additionally, when interacting genes were called on the basis of genomic proximity to the enhancers, we observed a similar trend (Supplementary Figure 11). These results indicate that LSD1’s role at enhancers is important for a precise transcription of their cognate genes.

To further corroborate if LSD1 and its catalytic activity are required for suppression of enhancer activity and associated genes, we utilized luciferase reporter assays in *Lsd1*-GT mESC. We selected 11 enhancers with two latent enhancers and at least one enhancer from each of poised, intermediate, and active enhancers that showed significant upregulation in *Lsd1*-GT mESC compared to WT-mESC and also showed an upregulation of the associated gene. 1.0 ~ 1.2 kb of the enhancer-containing regions were cloned downstream of the HSV-Thymidine Kinase promoter-driven firefly luciferase gene. *Lsd1*-GT mESC were transfected with a control plasmid or plasmids expressing either human LSD1 or the catalytically inactive LSD1-K661A mutant ^68^ along with the reporter plasmids. We found that LSD1’s catalytic activity is indeed required for suppression of all the active enhancers tested (*p* < 0.1, Student’s *t*-tests, Supplementary Figure 12), consistent with high levels of LSD1 at these enhancers in mESC. In contrast, one of the *Nanog* enhancers, which had not displayed a change upon the loss of LSD1 in mESC, was unaffected by LSD1 expression. We observed lower enhancer activities of the latent, poised, and intermediate enhancers compared to the active enhancers, indicating that our enhancer classification could accurately predict enhancer activity. However, we found it difficult to interpret the effect of LSD1 or its catalytic activity at these weak enhancers as they failed to enhance the activity of the promoter.

### Both mESC-specific and differentiation genes are upregulated upon the loss of LSD1 in undifferentiated mES cells

A previous study has implicated LSD1 in decommissioning of enhancers of pluripotency genes during differentiation of mES cells ^20^. However, the roles of LSD1 in undifferentiated mES cells remain elusive. The global upregulation of enhancers (Figure 4b), prompted us to investigate if the loss of LSD1 in undifferentiated mES cells affected the expression of pluripotency and/or differentiation genes. To this end, we first analyzed a published RNA-Seq datasets for mESC and epiblast stem cells (EpiSC), which were derived from mESC by treatment with Activin A and FGF2 for 4 days ^69^. We first selected the genes that showed a significant (*q* < 0.01, DESeq) and at least 5-fold change in expression during differentiation. This analysis yielded 710 induced and 745 downregulated (decommissioned) genes during the differentiation of mESC to EpiSC (Figure 6a). mESC and EpiSC represent two consecutive stages of embryonic development, namely pre-implantation and post-implantation, respectively. Therefore, these sets of upregulated and downregulated genes may represent the earliest transcriptional response of mES cells to the differentiation cue.

**Figure 6.**
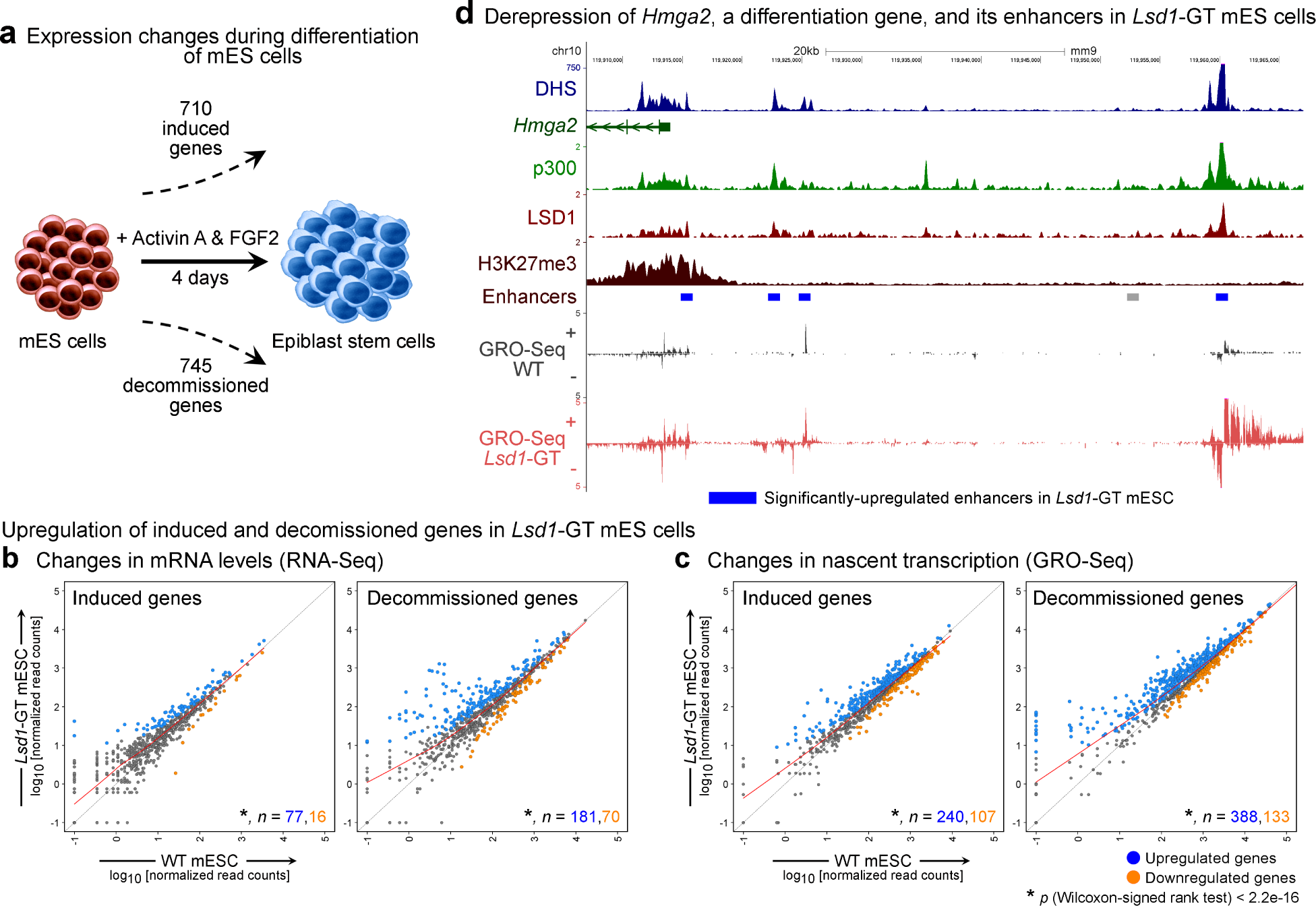
Both mESC-specific and differentiation genes are upregulated upon the loss of LSD1 in undifferentiated mES cells. (**a**) Schematic showing the number of significantly “induced” and “decommissioned” genes upon differentiation of mESC to epiblast stem cells with Activin A and FGF2 ^69^. (**b, c**) Scatter plots of mRNA levels (b) and levels of nascent transcription (c), as measured by RNA-Seq and GRO-Seq, respectively in WT and *Lsd1*-GT mESC. Number (*n*) of significantly-upregulated (*q* < 0.05) and -downregulated genes in each category are shown in blue and orange, respectively. Upon the loss of LSD1 in mESC, both groups of “induced” and “decommissioned” genes show a significant increase (*p* < 2.2e-16, Wilcoxon signed-rank test) in mRNA levels and nascent transcription. (**d**) Elevated transcription of *Hmga2*, a differentiation gene that plays an important role in exit of mES cells from the ground pluripotency state ^70^, and its nearby enhancers in *Lsd1*-GT mESC. Gray bar: Predicted enhancer. Blue bar: significantly-upregulated enhancer in *Lsd1*-GT mESC.

Compared to WT mESC, *Lsd1*-GT mESC displayed roughly equal number of genes being significant-upregulated or -downregulated (55% vs. 45%, 1493 upregulated and 1203 downregulated, respectively, *q* < 0.05). Quantitation of gene expression changes of the induced and the decommissioned genes, using our RNA-Seq and GRO-Seq datasets revealed that many of these genes are upregulated in undifferentiated *Lsd1*-GT mESC and both of these gene sets showed statistically-significant upregulation as group (*p* < 2.2e-16, Wilcoxon signed-rank test, Figure 6b, c). For example, *Hmga2* is a gene which is induced upon differentiation of mES cells and is required for the exit from naïve pluripotency ^70^. As shown in Figure 6d, LSD1 loss led to a marked increase in *Hmga2* expression which was also associated with increased H3K4 methylations, H3K27ac, and GRO-Seq signals at nearby enhancers. We have shown earlier that some pluripotency genes including *Pou5f1* (Figure 3b) and *mir290* (Supplementary Figure 10a) and their nearby enhancers were upregulated in *Lsd1*-GT mESC. Upregulation of both pluripotency and differentiation genes upon the loss of LSD1 in mES cells suggests that LSD1 does not instruct the fate of mES cells to a particular direction.

### LSD1 is required for suppression of inducible enhancers in terminally-differentiated neurons

We next sought to test if LSD1-mediated regulation of enhancers plays a role in gene expression program of terminally-differentiated cells using cortical neuron (CN) culture as a model. Using lentiviral delivery of two independent short hairpin RNAs (shRNAs) at 7 days *in vitro* (DIV), we knocked down (KD) LSD1 in primary cultures of mouse CN (Figure 7a). We employed BrU-Seq, a recently-developed nascent-RNA sequencing technique ^71^, that allows an accurate evaluation of any changes in active transcription of both mRNAs and eRNAs. At DIV 11, i.e. after 4 days of control or *Lsd1* shRNA delivery, neuronal cultures were treated with 5-bromouridine for 32 min, followed by enrichment of BrU-containing nascent transcripts using anti-BrdU beads and high-throughput sequencing. DESeq analysis indicated a significant misregulation of 1,500 genes (*q* < 0.05, 778 downregulated and 722 upregulated). Interestingly, many well-characterized activity-regulated genes (ARGs) ^43^, including *Arc, Fos, Fosb, Npas4, Egr1-4*, and *Nr4a1-3*, were among the most significantly-upregulated genes upon LSD1-KD in unstimulated neurons (Figure 7b). ARGs are expressed at low levels in resting neurons and are rapidly induced by depolarization of neurons via sensory inputs, thereby representing a stimulus-responsive gene regulatory program. Since products of ARGs play important roles in synaptic plasticity underlying cognitive development, learning and adaptive processes ^72-74^, we narrowed our focus on these inducible genes.

**Figure 7.**
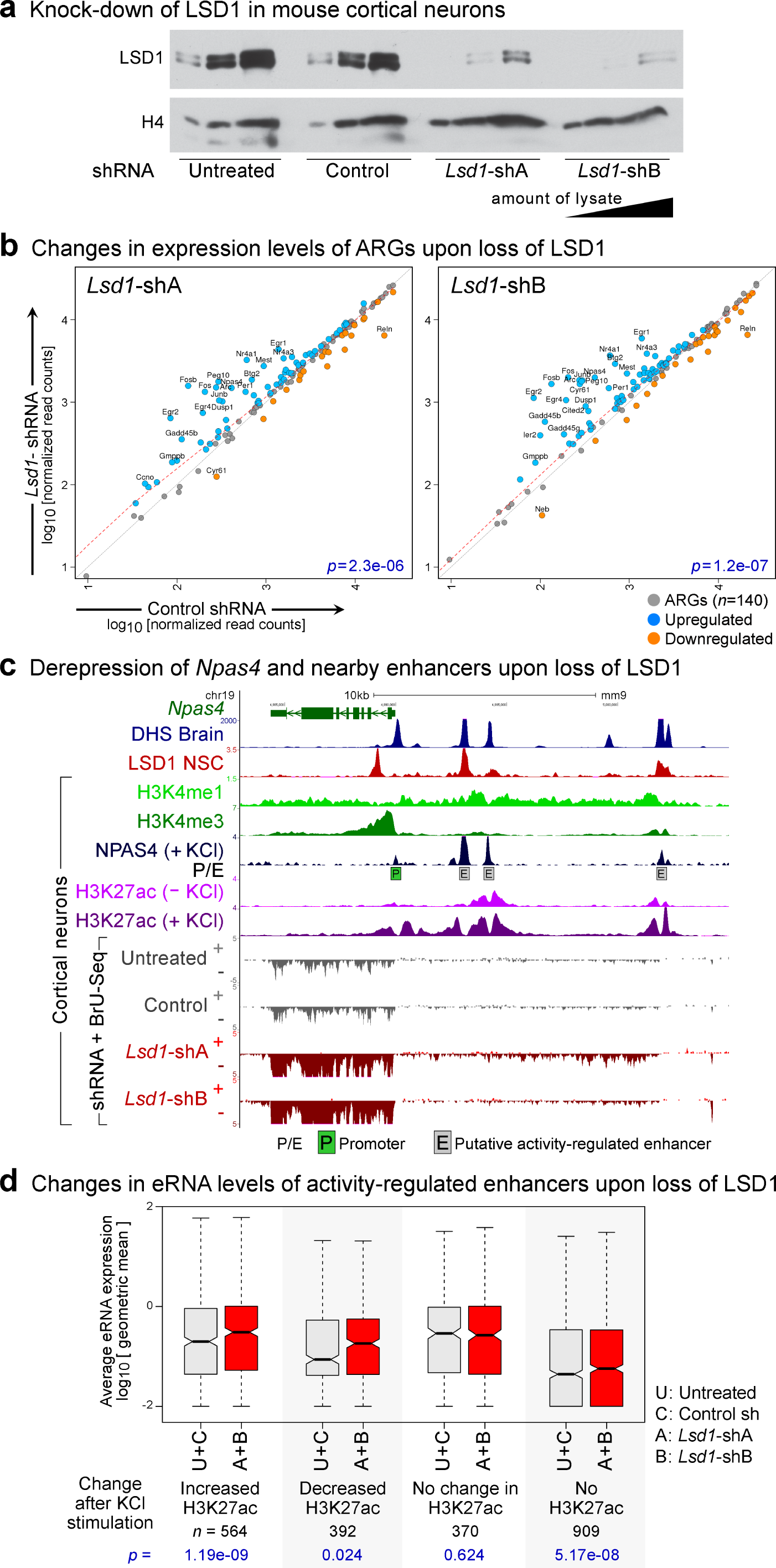
LSD1 is required for suppression of inducible enhancers in terminally-differentiated neurons. (**a**) Western blot analysis to confirm the knockdown (KD) of LSD1 in mouse cortical neurons (CN) at DIV 11, after 4 days of lentiviral-mediated delivery of either scrambled shRNA (control) or two independent *Lsd1* shRNAs (A and B). (**b**) Upregulation of activity-regulated genes (ARGs) in *Lsd1*-KD CN. Scatter plots of transcription levels of ARGs (*n*=140), from BrU-Seq analysis, in CN treated with either *Lsd1* shRNAs (Y-axis) or control shRNA (X-axis). Significantly-upregulated (*q*< 0.05, DESeq) and -downregulated ARGs are shown in blue and orange, respectively, and ARGs displaying greater than a 2-fold difference upon the loss of LSD1 are labeled with gene symbols. *P*-values (*p*) from Wilcoxon signed-rank tests are denoted in blue. (**c**) Aberrant induction of *Npas4*, an ARG, upon *Lsd1*-KD in resting CN. Boxed P: *Npas4* Promoter. Boxed E: Putative activity-regulated enhancers as evident from presence of DHS ^32^, high H3K4me1, low H3K4me3 ^75^, activity-dependent binding of NPAS4, and an increase in H3K27ac after KCl treatment ^76^. *Npas4* mRNA and eRNA are upregulated specifically in the *Lsd1*-KD neurons (red) (**d**) Increased eRNA levels at activity-regulated enhancers in *Lsd1*-KD CN. These enhancers have been previously divided into four groups based on the activity-induced changes in H3K27ac ^76^. Three groups of enhancers showed a significant increase in eRNA levels upon *Lsd1*-KD (red boxes) compared to control conditions (untreated CN or control shRNA-treated CN, gray boxes). A+B: Geometric mean of eRNA levels in CN treated with either *Lsd1* shRNAs A or B. U+C: Geometric mean of eRNA levels in control neurons. *P*-values (*p*) from Wilcoxon signed-rank tests are denoted in blue beneath each panel.

To evaluate this ARG upregulation on a genome-wide scale, we analyzed our previously published RNA-Seq data set ^75^ and identified 140 ARGs that were induced by KCl-mediated depolarization of CN in culture. Both of the two independent *Lsd1* shRNAs led to a spurious induction of many ARGs in the resting neurons (Figure 7b, Supplementary Figure 13), indicating that LSD1 suppresses premature induction of ARGs in CN. We next examined the *Npas4* locus to check if enhancer misregulation upon the loss of LSD1 could be involved in the premature induction of ARGs. We identified three putative enhancers upstream of the *Npas4* promoter based on DHS and H3K4me1 enrichment (Figure 7c). These three enhancers appear to respond to membrane depolarization, as they show activity-dependent increases in NPAS4 binding ^43^ and H3K27ac levels ^76^. These activity-regulated *Npas4* enhancers are bound by LSD1 in NSC ^52^. An increase in nascent transcription across these enhancers, concomitant with an increased *Npas4* expression, indicates that these enhancers are deregulated upon the loss of LSD1 (Figure 7c). A previous study had reported more than ten thousand putative activity-regulated enhancers based on increased CBP binding in response to membrane depolarization ^43^. Subsequent work categorized these candidate enhancers into four groups on the basis of activity-dependent changes in H3K27ac ^76^ (Figure 7d). The study found that only the enhancers that displayed activity dependent changes in H3K27ac were involved in promoting ARG transcription ^76^. Next, we investigated if the premature upregulation of ARGs in *Lsd1*-KD neurons was accompanied with misregulation of any of these activity-regulated enhancer groups. Analysis of BrU-Seq data revealed that loss of LSD1 did not have a significant impact on enhancers that do not display any activity-dependent changes in H3K27ac (Wilcoxon signed-ranked test, Figure 7d). However, *Lsd1*-KD led to a significant upregulation of eRNA levels at the enhancers that gain or lose H3K27ac upon KCl treatment (Figure 7d, Supplementary Figure 14a). Interestingly, the group of enhancers with no H3K27ac either before or after depolarization, and are presumably poised neuronal enhancers, also showed an upregulation upon the loss of LSD1 (Figure 7d, Supplementary Figure 14a). Similar to our earlier observations with RNA-Seq and Nuclear RNA-Seq in mESC, the eRNA signals with BrU-Seq were considerably lower than those with mESC GRO-Seq to obtain sufficiently-high statistical power for comparison; therefore, we aggregated eRNA signals from the two control and the two *Lsd1*-KD experimental groups for this analysis (Figure 7d, Supplementary Figure 14a). Similar trends were observed in analysis without grouping the samples (Supplementary Figure 14b). These data indicate that LSD1 is required for genome-wide suppression of premature enhancer activation in resting neurons.

Activation of ARGs upon *Lsd1*-KD could also be a result of extraneous activation of signaling pathways upstream of ARG induction. Extracellular signal-regulated kinases 1 and 2 (ERK1/2) are rapidly phosphorylated in response to a variety of extracellular stimuli, including membrane depolarization, and play critical roles to mediate the transcriptional response ^72, 77^. A lack of noticeable differences in phosphorylation levels of ERK1/2 upon *Lsd1*-KD (Supplementary Figure 14c), further support a direct role of LSD1 in suppression of activity-regulated enhancers and genes.

## Discussion

Early embryonic lethality of homozygous *Lsd1*-KO mice indicates an essential role of LSD1 in development ^78^. However, the roles of LSD1 in early embryogenesis have not been fully elucidated. A previous study has shown that the silencing of pluripotency genes during differentiation of mES cells is mediated by the decommissioning of pluripotency enhancers by LSD1 ^20^. This study employed Tranylcypromine (TCP), a pharmacological agent to block LSD1’s enzymatic activity ^20^. However, TCP also inhibits the H3K4 demethylase activity of LSD2 (aka KDM1B) ^79^, the paralog of LSD1, which is involved in regulation of transcriptional elongation ^80^. Thus, it remains unclear whether the observed impact of TCP treatment on enhancer dysregulation in mES cells was mediated by inhibition of either LSD1 or LSD2 or both. By employing genetic ablation of *Lsd1* in mES cells, we demonstrate that LSD1 suppresses the activity of a large fraction of primed enhancers, including the pluripotency enhancers and poised enhancers of differentiation genes. Notably, several of these key pluripotency enhancers and genes are already upregulated in undifferentiated *Lsd1*-deficient mESC (Figure. 3 and 6). Similar to our observations, another group had found that *Lsd1*-KD by siRNA led to an upregulation of several stem cell genes in undifferentiated mES cells ^81^. Loss of LSD1 in stem cells has been implicated in multiple differentiation defects, including de-differentiation of the pluripotent mESC state towards the totipotent 2-cell state ^82^, or the premature differentiation of human ES cells to endodermal and mesodermal lineages ^26^. These observations could be reconciled by our findings that LSD1 is required for suppressing both pluripotency genes and differentiation genes in mES cells, possibly through maintenance of proper enhancer activity.

We provide several lines of evidence that LSD1 plays an essential role in genome-wide homeostasis of primed enhancers. We show that recruitment of LSD1 correlates positively with levels of enhancer H3K4me2, H3K27ac and eRNA transcription (Figure 1) and this recruitment is specific to primed enhancers (Figure 2). Loss of LSD1 led to an upregulation of a large number of enhancers, as demonstrated by increased H3K4 methylation, H3K27ac (Figure 3), and eRNA transcription (Figure 4), concomitant with an upregulation of the associated genes (Figure 5). These results support the following model of LSD1-mediated homeostasis of the histone modification landscape during the life cycle of an enhancer. Binding of TF and subsequent recruitment of MLL3/4 ^14-16^ prime the enhancers with H3K4me1/me2, which attract LSD1 irrespective of whether the enhancers are destined to be either “active” or “poised” (Supplementary Figure 15). LSD1 then counteracts with MLL3/4 to maintain an optimal H3K4me levels. Enhancers with a relatively low H3K4me2 may represent early stages of priming by TFs and MLLs. It is possible that enhancers with low levels of H3K4me2 recruit little LSD1, which is not detectable by ChIP-Seq (Q1, Figure 1b). When gene expression needs to be increased, recruitment of additional factors and/or MLL3/4 may convert these less active enhancers to more active enhancers with higher H3K4 methylation and H3K27ac, which would then require higher levels of LSD1. LSD1’s recruitment might also serve as a surveillance mechanism to suppress ectopic installation of H3K4 methylation and spurious activation of enhancers.

During differentiation of mES cells, the pluripotency enhancers may be unprimed by the loss of ES-specific TFs followed by the loss of MLL3/4 and H3K4 methylation. Our model is not mutually exclusive to LSD1-mediated decommissioning of enhancers, as LSD1 could remove remnant H3K4me2/me1 to completely disengage the enhancer from active regulation. To further elucidate the mechanisms of decommissioning of pluripotency enhancers, it will be important to determine how early differentiation cues shift the balance of MLL3/4-mediated H3K4 methylation and LSD1-mediated demethylation.

Enrichment of H3K4me1 and depletion of H3K4me3 was the first combination of chromatin signatures to predict a large number of transcriptional enhancers in a mammalian genome ^10, 11^. More recent studies have shown H3K4me3 to be present at a subset of active enhancers ^83^ with a positive correlation between the H3K4me3/me1 ratio and enhancer transcription levels ^42^. We found that KDM5C and LSD1 can co-occupy enhancers in mES cells (Figure 4a). However, only the loss of LSD1, but not KDM5C, displayed significant changes in enhancer activity and gene expression, highlighting an essential and non-redundant role of LSD1 in mES cells. KDM5C has been implicated in both promotion of enhancer activity by generating H3K4me1 in mES cells ^22^, and suppression of over-activation of enhancers in breast cancer cells ^19^. Consistent with the former study, our analysis found a small reduction in eRNA levels in *Kdm5c*-KO mESC (Supplementary Figure 9). In addition to *Kdm5c*, other KDM5 family members, *Kdm5a* and *Kdm5b*, are also expressed in mES cells at similar levels and could possibly compensate for the loss of KDM5C. These observations suggest differential requirement of KDM5C in context of either different enhancers or different cell types.

Repeated “write-and-read” of histone modifications can form a feed-forward loop to allow TF-independent maintenance and propagation of chromatin status. Such models have been well established for the propagation of H3K27me3 ^84^ and H3K9me3 ^85^. Maintenance and propagation of H3K4me epigenetic memory across generations by LSD1 have been observed in worms ^86^ and mice ^87^. PHF21A (aka BHC80), another member of the LSD1 complex ^50, 58^, was the first known reader protein to recognize unmethylated H3K4 ^88^. This unique combination of an H3K4 demethylase and a reader of unmethylated H3K4, makes the LSD1-PHF21A complex an ideal candidate to exert a self-perpetuating “erase-and-read” mechanism. However, a positive correlation between LSD1 and H3K4me2 and LSD1’s absence at latent enhancers suggest that the role of LSD1 in maintaining the epigenetic memory could be limited to other genomic elements and warrants further investigation.

LSD1’s role in suppression of primed enhancers does not appear to be restricted to mES cells. Similar to our observations in mESC, our BrU-Seq analyses in post-mitotic neurons revealed that LSD1 suppresses premature activation of neuronal activity-regulated genes and enhancers (Figure 7). LSD1 has a neuron-specific isoform (neuroLSD1 or LSD1n) with four extra amino acids in the catalytic domain ^89^ and an altered substrate specificity which remains ambiguous ^49, 90^. Since our RNAi approach depleted both neuroLSD1 and canonical LSD1 in CN, it remains unclear if one or both of LSD1 isoforms mediate the suppression of activity-regulated enhancers and ARGs. Given that the genetic ablation of neuroLSD1 led to a downregulation of ARG expression ^49, 91^, it is more likely that the canonical LSD1, and not neuroLSD1, is involved in the suppression of activity-regulated enhancers. Loss-of-function *>LSD1/KDM1A* >mutations have been genetically associated with several neurodevelopmental conditions ^92-94^. These disorders could possibly be attributed to uncontrolled activation of activity-regulated genes and enhancers upon the loss of LSD1 and/or neuroLSD1. LSD1-mediated homeostasis of transcriptional enhancers, therefore, underlies various physiological processes including embryonic development and human cognitive function.

## Methods

See supplementary methods for details.

### Cell culture

*Lsd1*-WT and *Lsd1*-GT mESC have been described previously ^28^. *Kdm5c*-KO mESC were derived from the previously described mESC that carry the floxed *Kdm5c* allele ^75^ by Cre-mediated deletion of exons 11 and 12, which encode the enzymatic JmjC domain. mESC were grown on gelatin coated plates.

### Western blot analysis

mESC or CN were lysed in Laemmli sample buffer, sonicated, and subjected to SDS-PAGE. Western blot analyses were carried out using standard protocols using anti-H3K4me1 (ab8895, Abcam), anti-H3K4me2 (ab7766, Abcam), anti-H3K4me3 (ab8580, Abcam), anti-H3K27ac (39135, Active Motif), anti-KDM5C ^75^, anti-LSD1 (ab17721, Abcam) ^20^ and anti-phospho-ERK1/2 (4370, Cell Signaling Technology).

### *Lsd1* knockdown in mouse cortical neurons

Primary cultures of cortical neurons were carried out as described previously ^75^. *Lsd1*-KD in CN was achieved by lentiviral delivery of either scramble shRNA (SHC202, Sigma) or *Lsd1*-shRNAs (A: TRCN0000071375 and B: TRCN0000071376, Sigma) ^95^ on 7 days *in vitro* (DIV) and 5-Bromouridine incorporation was performed on DIV 11.

### ChIP-Seq

Antibodies used for chromatin immunopreciptation (ChIP) were anti-H3K4me1 (ab8895, Abcam and 07-436, EMD Millipore), anti-H3K4me2 (05-790, EMD Millipore), anti-H3K4me3 (04-745, EMD Millipore), anti-H3K27ac (39135, Active Motif), anti-HDAC1 (A300-713A, Bethyl Laboratories and sc-6298, Santa Cruz Biotechnology) and anti-KDM5C^75^. KDM5C ChIP-Seq experiments were performed as described previously ^75^. Other ChIP experiments were performed as described previously ^96^ with minor modifications.

### RNA-Seq and Nuclear RNA-Seq

RNA-Seq libraries have been described in detail previously ^97^. For sequencing of nuclear RNA, nuclei were isolated as described previously ^98^ with minor modifications. Libraries from rRNA-depleted RNA were prepared using Direct Ligation of Adapters to First-strand cDNA (DLAF) ^97^.

### Global Run-On

GRO was modified from the method described previously ^47, 98^. In addition to the presence of 0.2% IGEPAL CA-630, GRO on *Kdm5c* mESC, *Lsd1* mESC and CN were done in presence of 0.5%, 0.25% and 0.2% of *>N*-Lauroylsarcosine, respectively for 8 min at 30°C.

### BrU-Seq

Cortical neurons (DIV 11), after shRNA treatment for four days, were incubated with 2 mM 5-Bromouridine (850187, Sigma) for 32 min at 37°C. To reduce the number of steps for library preparation, we developed Direct Ligation of Adaptor to the 3’ end of RNA (DLAR), a method suitable for preparation of libraries for BrU-Seq.

All sequencing experiments were conducted in biological duplicates concurrently with different genotypes to minimize technical variations.

### Sequencing and Alignment

Multiplexed libraries were subjected to single-end sequencing on Illumina HiSeq 2000/2500 instruments using standard oligonucleotides designed for multiplexed paired-end sequencing, except that BrU-Seq indices were sequenced with DLAR_Index_Read:5’-CATAGGAAGAGCACACGTCTGAACTCCAGTCAC-3’.

ChIP-Seq reads were mapped to the mm9 genome using Bowtie1 (v1.1.2)^99^ allowing for up to two mismatches. PCR duplicates from ChIP-Seq reads were removed using samtools rmdup utility (v1.3) ^100^ and coverage along the genome was calculated using BEDTools (v2.25.0)^101^ after extending the ChIP-Seq reads to a total length of 180 bases. RNA-Seq libraries were mapped to the *mm9* genome and transcriptome using TopHat2 (v2.1.0)^102^ with Bowtie2 (v2.2.6) ^103^. For GRO-Seq and BrU-Seq, full length reads were first aligned using Bowtie1 or Tophat2, respectively. Adaptor sequences were trimmed from the unmapped reads using BBDuk utility ^104^ and reads were remapped and merged to the reads from the initial alignment. Only uniquely mapping reads were retained for further analysis and libraries were normalized to total number of non-mitochondrial and non-ribosomal reads.

### Analysis

MACS2 (v 2.1.0)^105^ was used to call DHS or ChIP-Seq peaks. For selection of candidate p300/DHS sites for enhancer prediction, we first scanned the genome for the strongest (with highest MACS2 signal) p300 or DHS site in a 1,250 base sliding window. When both p300 and DHS sites were present in the same window, p300 binding site was given higher precedence over any DHS sites. Intergenic p300/DHS sites were defined as sites that were outside of 1.25 kbp upstream to 3 kbp downstream of the genes. LSD1 has been shown to be involved in silencing of repetitive elements including endogenous retroviral elements (ERVs) ^28^. Therefore, to focus on prototypical enhancers in this study, we excluded p300/DHS sites with a low mappability (*M*_1_ < 0.75 and *M*_2_ < 0.75), where *M*_1_ and *M*_2_ indicate the fraction of uniquely mapping bases ^106^ within ± 500 and ± 100 bases, respectively, of the p300/DHS site. p300/DHS sites within the ENCODE blacklisted regions ^5^ were also excluded.

FeatureCounts ^107^ was used for calculating the number of reads overlapping various genomic features. Intersection analyses were done using BEDTools. DESeq (v1.22.1) ^63^ was used for normalization and differential gene expression analysis. 10-20 genes with exceptionally high expression and miRNAs and were excluded from further analysis. ChIP-Seq enrichment for H3K4me2, H3K4me3 and H3K27ac were normalized using MAnorm ^108^ with the fraction of reads aligning within the common peaks of WT and *Lsd1*-GT mESC samples from each replicate. ChIP-Seq coverage profiles from only one replicate, which did not require MAnorm normalization, were used in the browser snapshots in the figures. Prioritization of enhancer assignment is detailed in the supplemental information. Activity-regulated genes were identified as genes showing significant upregulation (*p* < 0.05, DESeq) in each of the two independent replicates of previously published RNA-Seq datasets from untreated and KCl-treated CN ^75^. Wilcoxon signed-rank tests were performed after log transformation of changes in expression or ChIP enrichment. The Perl scripts used for analyses are available upon request.

## Accession Numbers

Raw and processed sequence data files are available on the Gene Expression Omnibus (GEO) under accession GSE93952. The data can be accessed by the reviewers at: https://www.ncbi.nlm.nih.gov/geo/query/acc.cgi?token=ulchsiocxvojtaj&acc=GSE93952

## Acknowledgements

We thank Dr. Fulai Jin and Dr. Gary Hon at LICR, San Diego, for helpful discussions and guidance with bioinformatics. We are also grateful to Dr. Nima Mosammaparast at Washington University School of Medicine in St. Louis for the kind gift of WT and catalytically inactive *LSD1* >expression plasmids. The work was supported by grants from the LICR (to BR), the California Institute for Regenerative Medicine (RN2-00905, to BR), the University of Michigan Medical School (to SI), Cooley's Anemia Foundation Fellowship (to SI), NIH (NS089896, to SI), the Farrehi research fund (to SI), NSF Graduate Research Fellowship Program (DGE #1256260, to PMG), University of Michigan Career Training in Reproductive Biology (T32HD079342, to RSP), and the Eunice Kennedy Shriver National Institute of Child Health and Human Development DIR (HD008933, to TSM).

## Author contributions

SA and BR conceived the project and designed the experiments. SI and SA performed the RNA-Seq, Nuclear RNA-Seq, and GRO-Seq experiments. SA, EB, and TSM performed the ChIP-Seq experiments. SI, PMG, and SA designed and performed the BrU-Seq experiment. SA and RSP performed the luciferase reporter assays. YMN, PMG, and SA performed the western blot analyses. SA established the protocols and performed the computational analysis. SA and SI designed the analysis and wrote the manuscript. All authors approved and edited the manuscript.

